# Dimethyl sulfoxide primes induced pluripotent stem cells for more efficient nephron progenitor and kidney organoid differentiation

**DOI:** 10.1101/2025.02.07.637033

**Authors:** Helen Kearney, Aleksandra Rak-Raszewska, Adrián Seijas-Gamardo, Enrique Escarda-Castro, Paul Wieringa, Lorenzo Moroni, Carlos Mota

## Abstract

The field of human induced pluripotent stem cells (hiPSCs) has seen significant progress since the discovery of reprogramming somatic cells using the transcription factors Oct4, Sox2, Klf4, and c-Myc. hiPSCs are similar to embryonic stem cells in a primed state of pluripotency and has the potential to differentiate into any adult human cell type, offering a versatile tool for research and potential therapeutic applications. However, the efficiency of differentiation protocols for generating complex structures with multiple cell types, like kidney organoids, remains a challenge. This study investigates the impact of treating hiPSCs with a low-dose dimethyl sulfoxide to enhance kidney organoid differentiation using a well-established protocol from literature. We found that treating hiPSCs with 1-2% DMSO affects gene expression of pluripotent transcription factors, hiPSC colony morphology, and enhances the expression of key metanephric mesenchyme nephron progenitor marker, SIX2 after 9 days of kidney organoid differentiation. Our findings also suggest that DMSO treatment helps improve hiPSC differentiation protocol efficiency toward the development of tubular kidney organoids. Further research is needed to elucidate the mechanisms underlying these effects and to refine the differentiation process for potential *in vitro* research applications in biomedical research and drug development.

## 1 Introduction

In less than two decades since Yamanaka and colleagues first published a protocol to reprogram somatic cells back to a pluripotent state using key transcription factors such as Oct4, Sox2, Klf4, and c-Myc, the field of human induced pluripotent stem cells (hiPSCs) has undergone remarkable advancements [1, 2]. This Nobel-prize winning work has equipped researchers with the tools to induce pluripotency from any individual donor, directing differentiation towards any cell type in the body. Notably, hiPSCs also circumvents many ethical concerns associated with the use of embryonic stem cells, thereby opening new avenues in stem cell biology and its diverse applications [3]. hiPSCs are often considered to be similar to embryonic stem cells in a “primed state,” representing a more developmentally advanced pluripotent state similar to post-implantation epiblast cells [4, 5]. These cells can self-renew and differentiate into all three germ layers *in vitro*, but their ability to contribute to chimeras is limited. They grow as flat colonies and require FGF and Activin signalling for maintenance [6, 7]. These factors contribute to why hiPSCs are deemed to be in a more “primed state,” and serve as markers of pluripotency through colony morphology and the regulation of specific genes and cell surface proteins.

Recent research has highlighted key growth factors necessary to differentiate hiPSCs into kidney organoids, multicellular structures that closely mimic the fetal kidney development up to the second trimester [8-12]. Kidney organoids contain early-stage nephron structures, including glomeruli, proximal tubule, distal tubule, and collecting ducts, exhibiting a degree of functionality comparable to *in vivo* tissue. Kidney organoids hold significant promise for nephrotoxicity screening, potentially replacing animal models know for their poor predictive power [13]. However, current protocols for generating kidney organoids still face challenges such as limited efficiency, maturation, and the emergence of off-target cell populations with prolonged culture [14, 15]. Differentiation protocols require transitioning hiPSCs through early developmental stages, from primitive streak to intermediate mesoderm and nephron progenitors before forming kidney organoids [16]. Optimizing each stage of differentiation, particularly the starting hiPSC population, is crucial.

Previous studies show that treating human pluripotent stem cells (hPSCs) with a low concentration of dimethyl sulfoxide (DMSO) prior to directed differentiation increases differentiation across all germ layers. Chetty *et al*. show that DMSO prevents the phosphorylation of retinoblastoma, halting cells in the G1 phase of the cell cycle via alterations in PI3K pathway signalling, which they found regulates the early transitory states of hPSCs toward differentiation [17, 18]. Additionally, they discovered that genes involved in cytoskeletal dynamics, cilium assembly, and cell adhesion were particularly influenced by DMSO treatment. The integration of these signalling pathways regulates various developmental processes, including proliferation, differentiation, fate determination, apoptosis, migration, adhesion, and cell shape, ultimately impacting organogenesis. Other study have demonstrated that use of low concentrations of DMSO improved differentiation into various cell types [19]. However, limited knowledge is available on the effect of DMSO treatment on the differentiation efficiency of hiPSC-derived kidney organoids.

The primary objective of this study was to examine the effect of treating hiPSCs with a low dose of DMSO for 24 hours, and how it can enhance the efficiency of kidney organoid differentiation following a well established protocol from Morizane and collegues [20]. Firstly, we explored the potential impact DMSO has on the “primed state” of our hiPSCs, by measuring markers of pluripotency and examining hiPSC growth and colony morphology changes in real-time over 24hrs. Additionally, the downstream effects on kidney organoid differentiation was measured by quantifying the expression of proteins such as SIX2, a marker for metanephric mesenchyme (MM) nephron progenitor cells (NPC) [21], and other characteristic kidney organoid proteins markers, like podocalyxin (PODXL), megalin (LRP2), and GATA-binding protein 3 (GATA3) on the final day of differentiation [8-12].

## 2 Materials and Methods

### 2.1 Human induced pluripotent stem cell culture maintenance

hiPSC lines LUMC0031iCTRL08; LUMCi004-C (RRID:CVCL_ZA01), referred to hereafter as LUMC, HUMIMC101; TISSUi001-A (RRID:CVCL_WU56), referred to hereafter as H101, and HUMIMC107; TISSUi007-A (RRID:CVCL_WU62), referred to hereafter as H107, were maintained as colonies on 1% Geltrex (Gibco) coated Nunc™ cell culture treated plates (ThermoFisher) with mTeSRplus medium (STEMCELL Technologies). Cells were passaged once the colonies grew to the point where edges were rounded. Cell colonies were dissociated from culture wells using gentle cell dissociation reagent (STEMCELL Technologies) and washed with phosphate buffered saline (PBS) (Gibco) before scraping from surface using a sterile cell lifter. Colonies were gently broken up by pipetting and reseeded at the desired splitting ratio, normally 1:10.

### 2.2 Seeding hiPSCs for differentiation

Cells were differentiated following a previously published protocol for differentiating hiPSC toward MM NPC and kidney organoid formation, and optimized for the LUMC hiPSCs following guidelines in Morizane *et al*. [20]. Briefly, 1mL Accutase (STEMCELL Technologies) was added to a well of hiPSC and incubated at 37 °C for 10min. A single cell suspension of hiPSCs resuspended in mTeSRplus medium supplemented with 10 μM Y-27632 dihydrochloride (Tocris) and seeded onto cell culture treated plates previously coated with 1% geltrex (ThermoFisher) solution according to manufacturer’s guidelines at a specific cell density for each hiPSC line used: 1 × 10^4^ cell/cm^2^ for LUMC, 9 × 10^3^ cell/cm^2^ for HUMIMC101, and 7 x10^3^ cell/cm^2^ for HUMIMC107. The following day, media was removed and mTeSRplus media was added for further 24hr. On the third day of culture, medium was changed to mTeSRplus medium supplemented with DMSO; none (negative control), 1%, or 2% v/v for 24 hrs.

### 2.3 Pluripotency cell surface markers by flow cytometry

Medium supplemented with DMSO was removed from hiPSCs cultured in 6-well plate after 24 hr and wells were washed once with PBS. 1mL Accutase (STEMCELL Technologies) was added to each well and incubated at 37 °C for 10 min. Cell solution was gently pipetted to obtain a single cell suspension. A panel of fluorophore-conjugated pluripotency marker antibodies, TRA-1-81, TRA-1-60, SSEA3 and SSEA4 (BD Bioscience) was diluted in flow cytometry stain buffer (BD Bioscience) following suppliers’ guidelines to assess pluripotency (Table S1). An amount of 1 x10^6^ cells were incubated with each antibody solution for 30 min in the dark at 4 °C. Cells were centrifuged at 300 g for 5 min and re-suspended in flow cytometry stain buffer. Flow cytometry was performed on BD Accuri C6 flow cytometer. Compensation parameters were assessed for each antibody and used to improve accuracy of final readout using FloJo analysis software.

### 2.4 Pluripotency intracellular markers by flow cytometry

Medium supplemented with DMSO was removed from hiPSCs cultured in 6-well plate after 24hr and wells washed once with PBS. 1 mL Accutase (STEMCELL Technologies) was added to each well and incubated for 10 min at 37 °C. Cell solution was gently pipetted to obtain a single cell suspension. Accutase was deactivated with an equal part of DMEM medium and cells centrifuged at 300 g for 5 min. Supernatant removed and cell pellet re-suspended in 4% PFA and incubated for 30 min at 4 °C. Cells centrifuged at 300 g for 5 min and resuspended in permeabilization buffer (0.1% Triton X-100 in PBS) and incubated for 10 min at room temperature. Fluorescently conjugated antibodies (SOX2-PE and OCT3/4-AF647), were diluted in flow cytometry stain buffer (Table S1). 1 × 10^6^ cells were incubated with each antibody solution for 30 min in the dark at 4 °C. Cells were centrifuged at 300 g for 5 min and re-suspended in flow cytometry stain buffer. Flow cytometry was performed on BD Accuri C6 flow cytometer. Compensation parameters were assessed for each antibody and used to improve accuracy of final readout using FloJo analysis software.

### 2.5 G1 synchronisation analysis

Medium supplemented with DMSO was removed from hiPSCs cultured in 6-well plate after 24 hr and wells were washed once with PBS. 1 mL Accutase (STEMCELL Technologies) was added to each well and incubated at 37 °C for 10 min. Cell solution was gently pipetted to obtain a single cell suspension. Samples were centrifuged at 300 g for 5 min. Supernatant was removed and the cell pellet was re-suspended in 70% ethanol (diluted in MilliQ water). Cell suspension was incubated overnight at 4 °C followed by centrifugation at 300 g for 5 min. Supernatant was removed and the cell pellet was re-suspended in flow cytometry stain buffer supplemented with 50 µg/mL propidium iodide solution and 200 µg/mL RNAse and incubate for 3 hrs at 4 °C to stain DNA and remove RNA in each sample. Flow cytometry was performed on BD Accuri C6 flow cytometer. Data analysed using FloJo analysis software.

### 2.6 Kidney organoid differentiation

The protocol previously established by Morizane *et al*. for kidney organoid differentiation [20] was followed in this study. Firstly, basal media composed of Advanced RPMI (Gibco) and 1% GlutaMax (Gibco) was supplemented with CHIR99021 (Tocris) and 5ng/mL recombinant human noggin (Peprotech), was added to hiPSCs following the 24hr DMSO treatment previously described. Each different hiPSC line required a different concentration of CHIR99021; LUMC – 8µM CHIR99021, HUMIMC101 – 10µM CHIR99021, and HUMIMC107 - 12µM CHIR99021. Medium supplemented with CHIR99021 and noggin was refreshed after 2 days. On day 4, cell colony morphology was assessed. At this point colonies had condensed into tightly packed colonies with smooth edges. Medium composed of Advanced RPMI supplemented with 1% GlutaMax and 10ng/mL Activin A (Miltenyi Biotech), was added to cells for 3 days. On day 7 of differentiation, spent medium was removed and medium composed of Advanced RPMI supplemented with 1% GlutaMax and 10 ng/mL FGF9 (STEMCELL Technologies), was added to cells. By day 9 of the differentiation, vesicles were visible under brightfield microscopy indicating a successful MM NPC differentiation. Medium was refreshed every 2-3 days of culture. On day 14 of differentiation medium composed of Advanced RPMI supplemented with 1% GlutaMax was added to the culture and refreshed every 2-3 days. Kidney organoids with defined characteristic tubules visible under brightfield microscope were obtained on day 21 of culture. The number of individual organoid structures in each well of the 96 well plate was manually counted by eye.

### 2.7 Generating LUMC-GFP+ hiPSCs

For the genetic modification of the LUMC hiPSCs the super piggyBac transposase (Hera BioLabs) and LipofectamineTM Stem Cell Transfection Reagent (Invitrogen) were used following instructions from the manufacturers. To generate the LUMC-GFP+ line, LUMC hiPSCs were passaged as single cells using Accutase on a Geltrex coated 24-well plate to a density of 75 × 10^3^ cells/well. Cells were maintained with mTeSR Plus media supplemented with 10 µM of Y-27632 (STEMCELL Technologies) for 1 day. On the next day 50 µL of Opti-MEM media containing 1 µL of Lipofectamine, 500 ng of the PiggyBac plasmid and 500 ng of the plasmid of interest (pSH231-EF1-GFP-HYGRO) was prepared. pSH231-EF1-GFP-HYGRO was kindly provided by Raymond Monnat & Michael Phelps (Addgene plasmid # 115144 ; http://n2t.net/addgene:115144 ; RRID:Addgene_115144) [22]. This suspension was then added to each well containing the hiPSCs with fresh mTeSR Plus media and incubated overnight in cell culture conditions. The next day the media was refreshed to remove the dead cells and 200 µg/mL of Hygromycin (STEMCELL Technologies) for positive selection of the transfected cells. These cells were then further passaged as single cells to generate colonies that later were hand-picked for achieving a homogenous cell population. Transfected LUMC-GFP+ cells were then assessed for pluripotency using SOX2 and OCT4 markers (Figure S2). hiPSC maintenance and differentiation was carried out following the same protocol as non-transfected LUMCs.

### 2.8 Immunofluorescent staining hiPSCs

All medium was removed from each well of 96-well plate and washed with PBS and cells fixed with 4% paraformaldehyde (Sigma) at 4 °C overnight. Samples were washed with PBS and stored for up to 4 months at 4 °C. Fixed plates were equilibrated at room temperature for 30 min to allow geltrex to fully crosslink before proceeding with staining protocol. Permeabilisation buffer composed of 0.1% v/v Triton-X (Sigma) was added to fixed cell culture for 10 min at room temperature. Blocking buffer composed by 3% w/v bovine serum albumin (BSA) (Sigma), was added to fixed cell culture for 1 hr at room temperature. Primary antibody solutions were prepared by diluting antibodies/stains for specific proteins at desired concentration (Table S1) in 1.5% BSA and incubated with fixed samples overnight at 4 °C. Plates were removed from the fridge and allow to warm up to room temp for 30 min. Samples were washed three times with PBS at room temperature. Secondary antibody solutions were prepared by diluting fluorophore-conjugated antibodies including DAPI at desired concentration (Table S1) in 1.5% BSA and incubated with fixed samples for 1 hr at 4 °C. Samples were washed three times with PBS at room temperature. All solutions made with 1X PBS were filtered through 0.2 µm syringe filter unit (Millipore) before the antibodies were added.

### 2.9 Immunofluorescent staining nephron progenitors and kidney organoids

All medium was removed from each well of 96-well plate and washed with PBS and cells/organoids were fixed with 4% paraformaldehyde (Sigma) at 4 °C overnight. Samples were washed with PBS and stored for up to 4 months at 4 °C. Fixed plates were allowed come to room temperature for 30 min, permeabilized with buffer composed of 0.1% v/v Triton-X (Sigma) for 15 min at room temperature. Blocking buffer composed by 0.3% v/v Triton-X, 0.05% Tween 20 (Sigma) and 3% bovine serum albumin (BSA) (Sigma), was added to fixed cell culture for 2 hr at room temperature. Primary antibody solutions were prepared by diluting antibodies/stains for specific proteins at desired concentration (Table S1) in 1.5% BSA and incubated with fixed cell culture overnight at 4 °C. Plates were brough to room temp for 30 min and washed three times with permeabilisation buffer for 30 min. Secondary antibody solutions were prepared by diluting fluorophore-conjugated antibodies including DAPI at desired concentration (Table S1) in 1.5% BSA and incubated with fixed samples overnight at 4 °C. Plates were placed at room temp for 30 min and washed three times with permeabilisation buffer for 30 min.

### 2.10 Image acquisition and processing

Images were acquired on automated inverted Nikon Ti-E microscope, equipped with a Lumencor Spectra light source, an Andor Zyla 5.5 sCMOS camera, and an MCL NANO Z200-N TI z-stage (Figure 1, 4, 5 and S2), and the Leica TCS SP8 inverted laser scanning confocal microscope (Leica Microsystems) (Figure 3). Live cell imaging was performed on an automated inverted Nikon Ti-E microscope, equipped with a Lumencor Spectra X light source, Photometrics Prime 95B sCMOS camera, an MCL NANO Z500-N TI z-stage, and a Okolab incubator (37 °C, 5% CO_2_) (Figure 2). Images were processed using NIS software (Nikon) and Fiji ImageJ.

**Figure 1.**
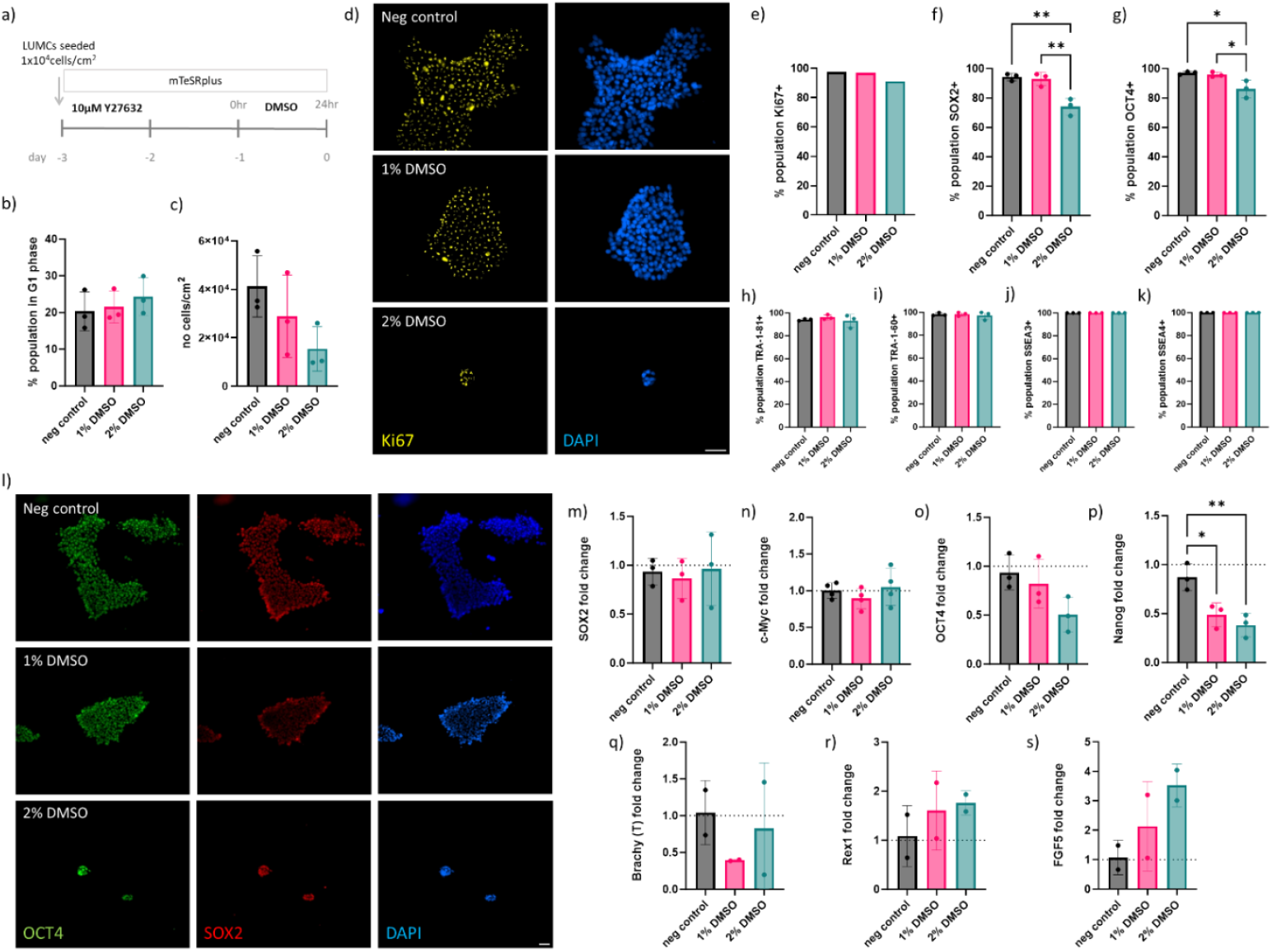
Pluripotency marker expression and cell cycle analysis: a) Schematic of cell seeding and 3 days of hiPSC culture including 24 hr DMSO treatment before commencing differentiation protocol. Characterization and quantifications performed at endpoint - day 0. b) The percentage of cells in G1 phase, and c) the number of hiPSCs counted/cm^2^ (N=3). d) Immunofluorescent images of hiPSC colonies expressing Ki67 and stained with DAPI, scale bar – 50 µm. e) The percentage hiPSCs expressing proliferation marker Ki67 (N=1). F-g) The percentage hiPSCs expressing pluripotent transcription factors SOX2 and OCT4 measured by flow cytometry (N=3). H-k) The percentage hiPSCs expressing pluripotent cell surface markers; TRA-1-81, TRA-1-60, SSEA3 and SSEA4 measured by flow cytometry (N=3). K) Immunofluorescent images of hiPSC colonies expressing OCT4 and SOX2 and stained with DAPI, scale bar – 50 µm. Gene expression analysis by qPCR of l-o) transcription factors for pluripotency; Nanog, c-Myc, SOX2, and OCT4 (N=4), p) Brachy (T) marker for mesoderm differentiation (N=4), q) Rex1 marker for naïve pluripotent stem cells (N=4), and r) FGF5 marker for primed pluripotent stem cells (N=4). Fo– p) - r), each data point refers to pooled cDNA from 2 biological replicates.

**Figure 2.**
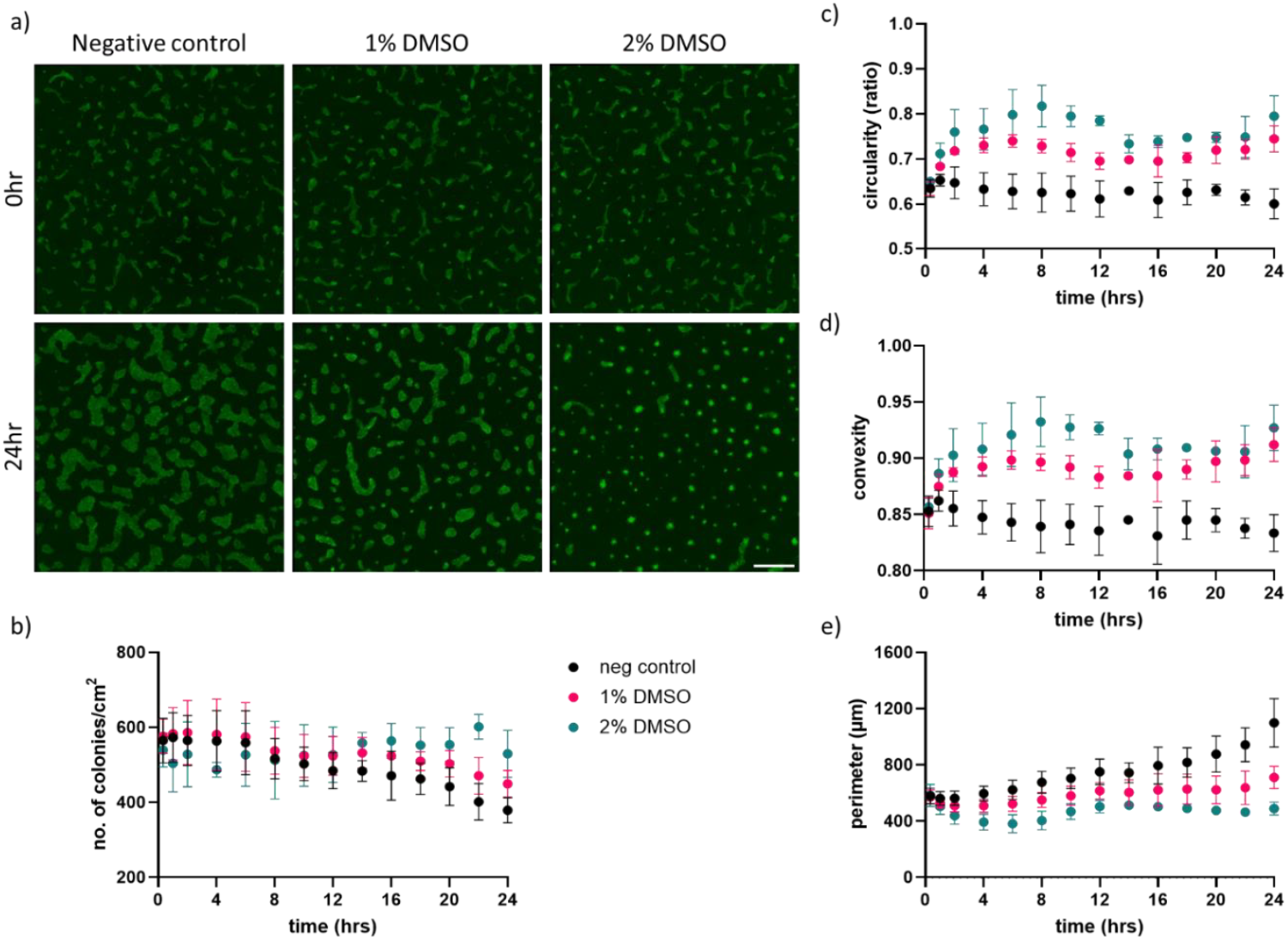
Parametric analysis of LUMC-GFP+ hiPSC colony morphology. a) Fluorescence images of LUMC-GFP+ hiPSC colonies at the beginning (0 hr, upper panel) and end (24 hrs, lower panel) of the exposure to DMSO. Scale bar of 1000 µm. Real-time measurements of b) number of colonies/cm^2^, c) circularity, d) convexity and e) perimeter over 24hr DMSO treatment (N=3).

**Figure 3.**
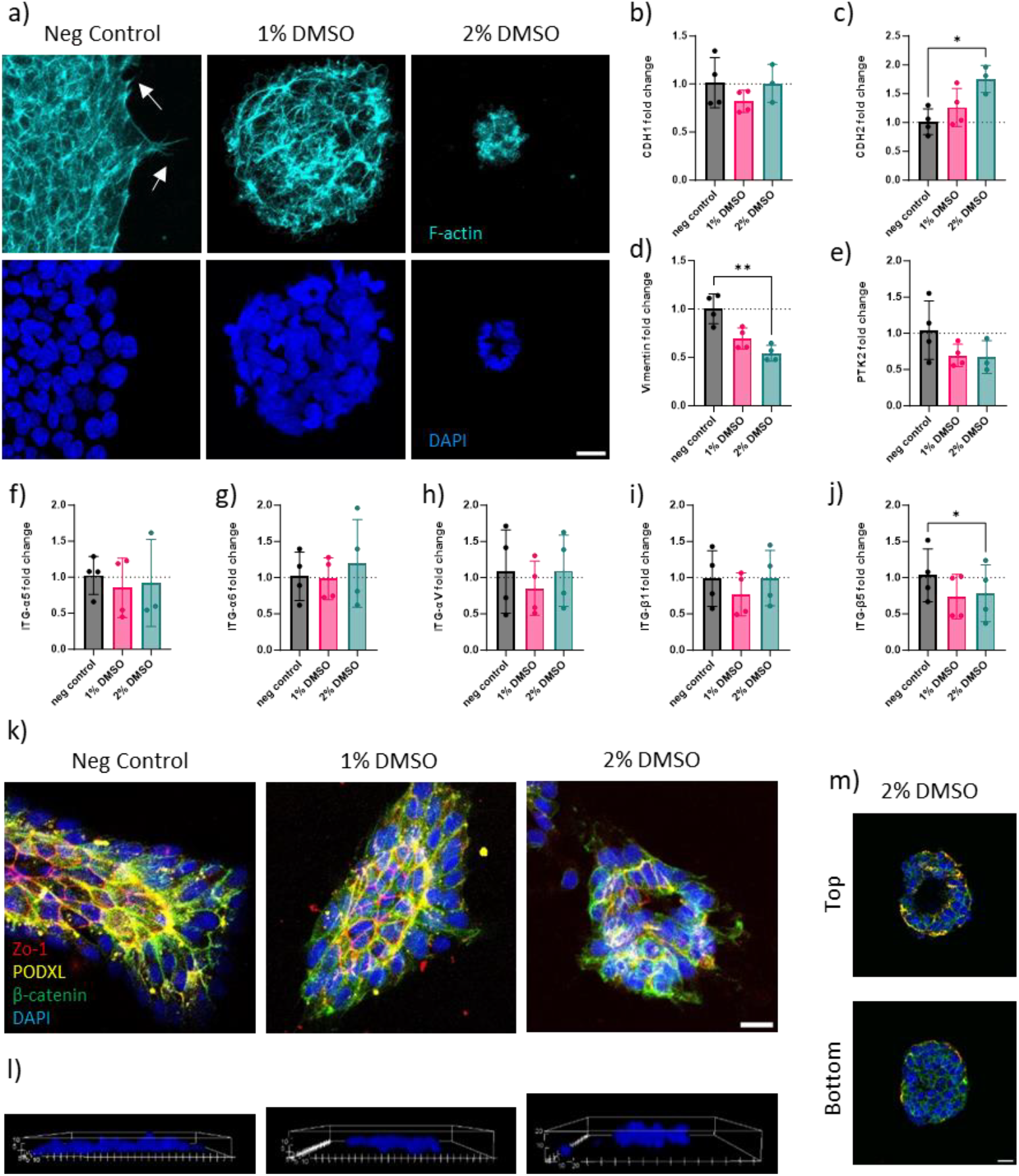
Changes in LUMC colony structural organisation following DMSO treatment. a) Immunofluorescence images of F-actin cytoskeleton stained with Phallodin (cyan) and DAPI (bue). Cellular protrusion indicative of focal adhesions indicated with white arrows. Gene expression analysis of b) E-cadherin (CDH1), c) N-Cadherin (CDH2), and d) Vimentin. Gene expression analysis of e) focal adhesion-associated protein kinase (PTK2), and f) integrin alpha-5 (ITG-α5), g) integrin alpha-6 (ITG-α6), h) integrin alpha-V (ITG-αV), i) integrin beta-1 (ITG-β1), j) integrin beta-5 (ITG-β5) (n=4). k) Immunofluorescence images of hiPSC colonies stained for β-catenin (green), ZO-1 (red), and podocalyxin (yellow) acquired on confocal and shown as 2D maximum intensity projection of an acquired z-stack and l) 3D rendered images of a representative colony showing height of colonies in z-plane. m) Single stack image from Z-stack from upper section (top) and lower section (bottom) of a representative hiPSC colony treated with 2% DMSO. Scale bar of 20 µm applicable to all images.

**Figure 4.**
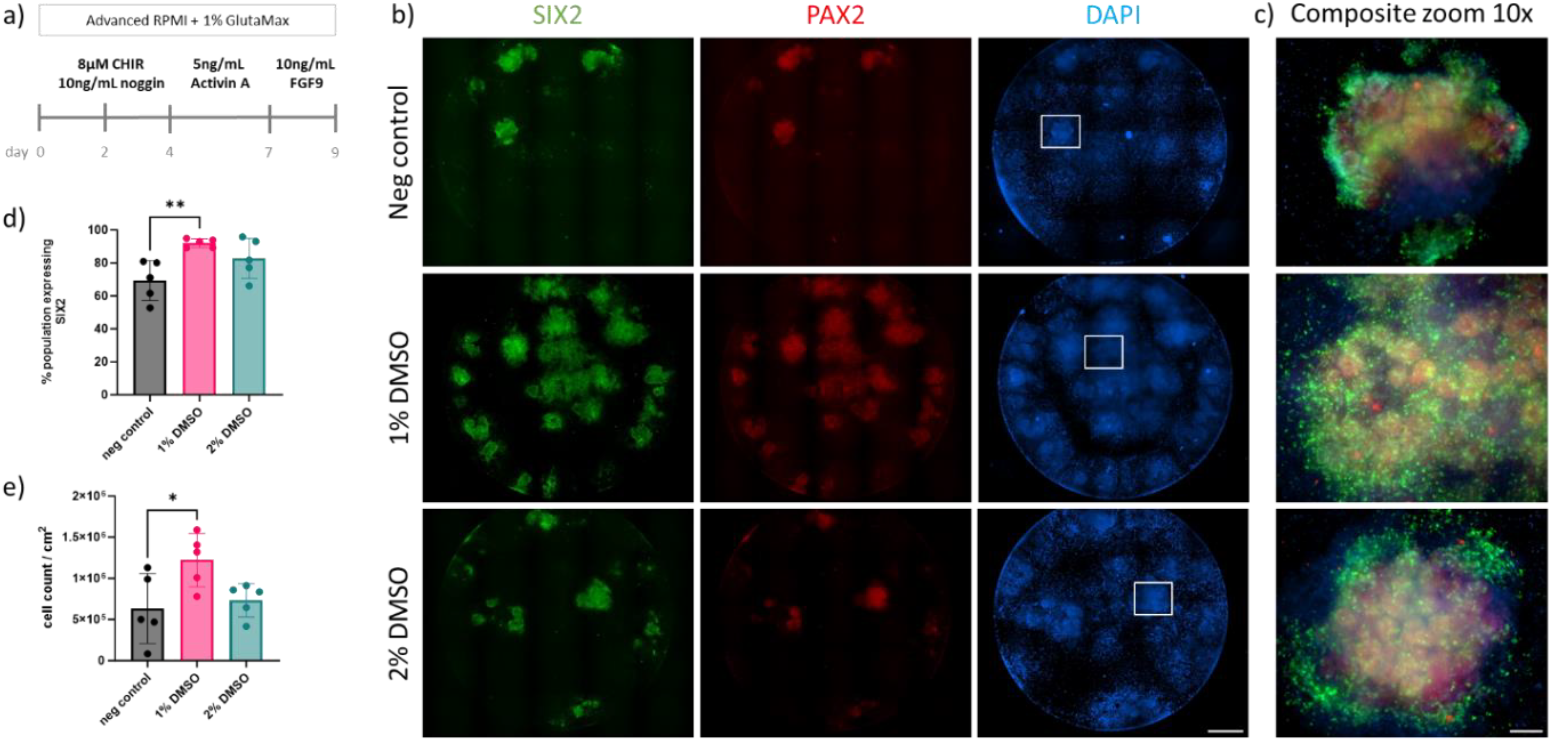
LUMC-derived metanephric mesenchyme nephron progenitor cells on day 9 of kidney organoid differentiation. a) schematic of the first 9 days of kidney organoid differentiation protocol up to the stage when nephron progenitors emerge. b) Immunofluorescence images of differentiated hiPSCs stained for MM markers; SIX2 (green) and PAX2 (red). Scale bar – 1000 µm. c) Composite zoom 10x images, scale bar – 100 µm. d) Percentage of SIX2+ cells at day 9, measured by flow cytometry (N=5). e) The number of cells/cm^2^ counted following dissociation using a manual haemocytometer (N=5).

**Figure 5.**
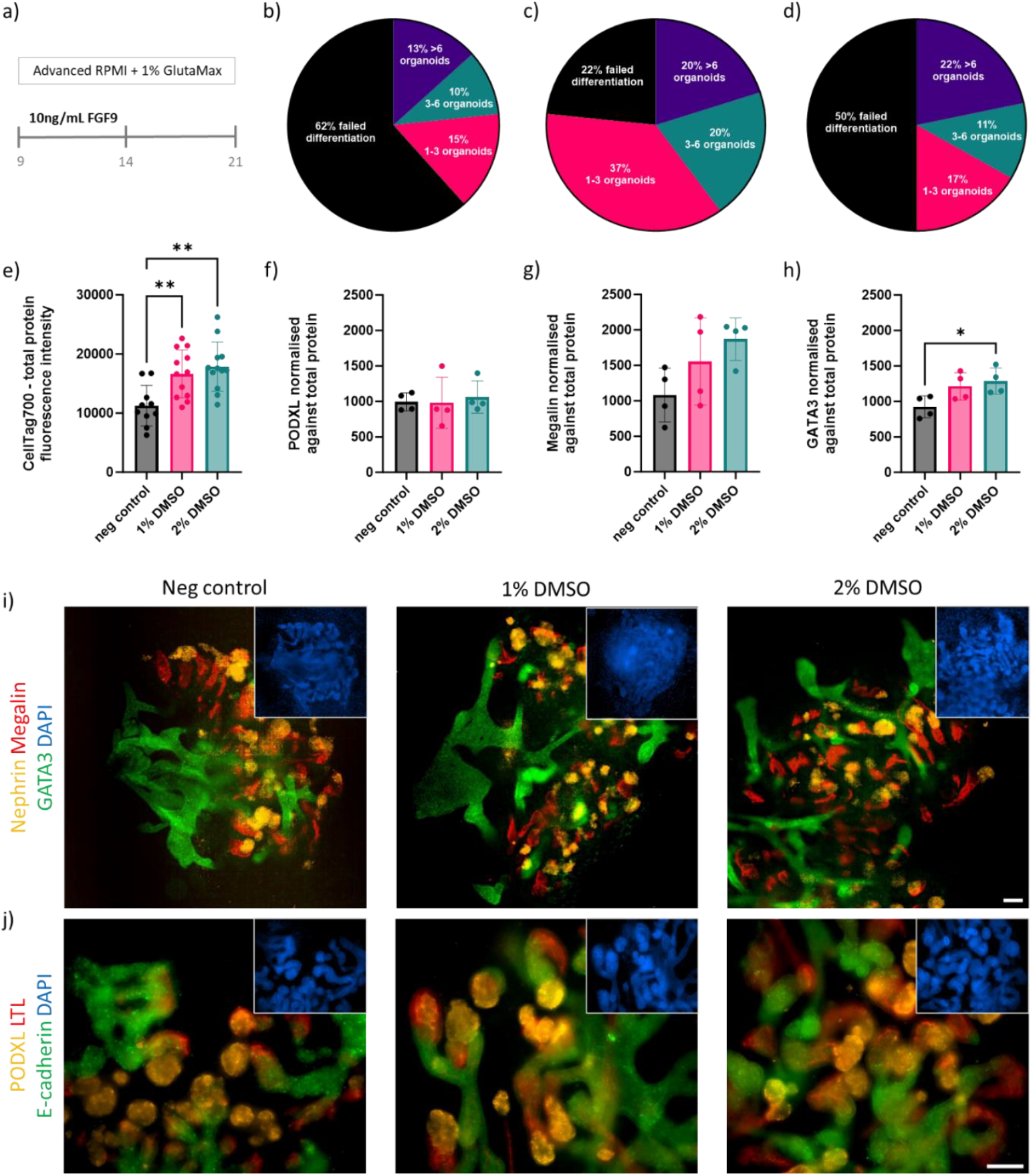
Characteristic LUMC-derived kidney organoid protein expression images and differentiation efficiency quantification on day 21 of the protocol. a) Schematic of the 11 days of differentiation from SIX2+_nephron progenitors to kidney organoid up to day 21. b-d) Percentage of wells (n=60) with successfully differentiated kidney organoid colonies developing; b – no treatment, c – 1% DMSO, d – 2% DMSO. e) Total fluorescent signal measured in each well (n=5) and used to calculate normalised signal of characteristic proteins found in different sections of the developing nephron shown in: f) podocalyxin – glomerulus, g) megalin - proximal tubule, h) and GATA3 - distal tubule/collecting duct. Representative immunofluorescence images of day 21 kidney organoids showing a general overview of each well with characteristic proteins found in different sections of the developing nephron: i) nephrin – glomerulus (yellow), megalin - proximal tubule (red), and GATA3 - distal tubule and collecting duct (green). j) podocalyxin – glomerulus (yellow), lotus tetragonolobus lectin (LTL) – proximal tubule (green), and E-cadherin – distal tubule (red). Scalebar of 100 µm valid for all images.

### 2.11 Quantitative PCR

All medium was removed from each well of 6-well plate wells. Plates were placed on ice and ice-cold PBS was added to each well for 5-10 min to dissolve Geltrex [23]. Cells were pipetted gently to resuspend, transferred to Eppendorfs and centrifuged at 750 g for 5 min at 4 °C. Cells were washed twice in ice cold PBS to remove all Geltrex and cell debris. The final cell pellet was dissociated and lysed using Trizol (Invitrogen) and stored at -80 °C until further use. Samples were thawed on ice and 20% v/v chloroform (Sigma) was added to lysed cell solution in trizol and centrifuged at 12000 rpm for 15 min at 4 °C to separate phases. Upper aqueous phase containing RNA was transferred to a new Eppendorf tube and precipitated in isopropanol (Fisher BioReagents) for 2 hr at - 20 °C. RNA precipitates were centrifuged at 12000 rpm for 30 min at 4°C. RNA pellets were washed with ice-cold 75% v/v ethanol twice and centrifuged at 7500 rpm for 10 min at 4 °C. The final pellet was left to airdry for 10 min at room temperature and diluted in nuclease-free water (Qiagen). The RNA concentration was measured with the BioDrop μLITE device. RNA was stored at -80 °C until further use. The iScript cDNA sysnthesis kit (BioRad) was used to synthesize a quantitative amount of cDNA by reverse transcription. Briefly a mixture of 1 μg RNA template, 1 μL iScript reverse transcriptase, 1x iScript reaction mix, was prepared with nuclease free water up to 20 μL final volume. This solution was mixed on PeqStar thermal cycler (PeqLab) following iScript kit instructions (25 °C for 5 min, 46 °C for 20 min, 95 °C for 1 min, and finally, hold at 4 °C). Double stranded cDNA was stored at -20 °C. For qPCR, cDNA was added to the wells containing 0.5 µM validated gene specific forward primer and reverse primers (Table S2) and iQ SYBR green supermix (BioRad) to obtain a final volume of 10 µL. Plates were sealed and incubated on BioRad CFX96 following iQ SYBR green supermix kit instructions; (95 °C for 1 min) x1, (95 °C for 15 s, 60 °C for 45 s) x 39, (95 °C for 15 s) x1. All genes of interest were normalised against verified stable mitochondrial housekeeper gene nuclear encoded (ATP5PB).

### 2.12 SIX2 protein quantification by flow cytometry

Cells were detached from each well of 6-well plate using Accutase for 15 min at 37 °C, followed by washing and straining to remove debris and cell aggregates. Cells were counted, centrifuged, and PBS was removed. One-third of each condition was stained with fixable live/dead stain (1:1000 in PBS) in 50 µL for 30 min on ice, protected from light. After adding 200 µL of FACS buffer (0.5% BSA in PBS), cells were centrifuged at 300 g for 3 min. The stain was removed, and cells were fixed with 2% PFA for 20 min at 4 °C. Cells were then washed with 300 µL FACS buffer and centrifuged again at 300g for 3 min. The fixative was removed, and cells were stored in FACS buffer at 4 °C until ready to stain. For staining, cells were centrifuged at 500 g for 3 min, treated with 50 µL of 0.1% Triton X for 15 min on ice, and centrifuged again at 500 g for 3 min. Cells were then centrifuged and incubated with 50 µL primary antibody solution (SIX2, 1:500, Proteintech) for 30 min on ice. After adding 1000 µL of PBS, cells were centrifuged again and incubated with 50 µL secondary antibody solution (Donkey anti-rabbit AlexaFluor488, 1:2500, Invitrogen) in 1.5% serum solution for 15 min on ice. The cells were washed with 1000 µL PBS, centrifuged, and resuspended in 200 µL 0.5% BSA. Beads were vortexed, and 10 µL of each sample (fluorophore only) was taken. Samples were incubated with 50 µL antibody solution for 30 min at room temperature in the dark, washed with 1000 µL FACS buffer, centrifuged, and resuspended in 200 µL 0.5% BSA. Each sample was then analysed using spectral flow cytometer Cytek Aurora (Cytek Biosciences), measuring AF 405 to detect dead cells (Live/Dead™ Fixable Aqua Dead Cell Stain Kit, ThermoFisher) to exclude them from analysis and AF488 to identify SIX2+ cell population. Unstained cells and live/dead stain cells were used to establish fluorescence background and define gates that would select single and live cells before analysing SIX2 stained samples. We analysed over 30 × 10^3^K cells per sample ensuring the live cell count was above 10 × 10^3^. Data analysis was performed using flow cytometer software, where percentage of SIX2+ population was given per at least 10 x10^3^ live cells counted. Further analysis was performed using GraphPad Prism (see below).

### 2.13 In-cell western

Selected wells that contained kidney organoids, cultured in Greiner 96 well plate, were washed with PBS to remove all media and fixed with 4% paraformaldehyde (Sigma) at 4 °C overnight. Samples were washed with PBS and stored for up to 4 months at 4 °C. Fixed plates were allowed to equilibrate to room temperature for 30 min to allow geltrex to fully crosslink before proceeding with staining protocol. Permeabilisation buffer composed of 0.1% v/v Triton-X (Sigma) was added to fixed cell culture for 15 min gently rocking at room temperature. Blocking buffer composed of 0.3% v/v Triton-X, 0.05% Tween 20 (Sigma) and 3% bovine serum albumin (BSA) (Sigma), was added to fixed cell culture for 2 hr gently rocking at room temperature. Primary antibody solutions were prepared by diluting antibodies/stains for specific proteins at desired concentration (Table S1) in 1.5% BSA and incubated with fixed cell culture for 24 hr at 4 °C. Plates were equilibrated to room temp for 30 min and washed three times with permeabilisation buffer for 30 min at room temperature. Near infrared secondary antibody and CellTag700 total protein stain were added at desired concentration (Table S1) to 1.5% BSA and incubated with fixed cell culture overnight at 4 °C. After being placed at room temperature for 30 min, the samples were washed three times with 0.05% v/v tween-20 (Sigma) diluted in PBS for 15 min. LICOR Odyssey CLX was used to acquire near infrared (NIR) fluorescent signal at 4 mm offset height. Signal intensity was quantified using LICOR Image Studio Ver 5.0 software. In each plate one well was incubated with each secondary antibody only and used to measure background signal. Total protein staining signal was used to normalize the staining signal for each of the proteins of interest.

### 2.14. Statistical analysis

Statistical analysis was performed using GraphPad Prism8 (version 8.2.0) software. One-way ANOVA with Tukey’s multiple comparisons was used for all qPCR results, flow cytometry, cell count and in-cell western protein quantification. A p-value smaller than 0.03 was considered statistically significant (denoted with *p < 0.05, **p < 0.01, ***p < 0.001, ****p< 0.0001). Results are shown as mean ± standard error of mean.

## 3. Results

### 3.1 Pluripotency and differentiation potential

To investigate the effect of DMSO on the hiPSCs line used, quantitative and qualitative assessment of key markers was performed three days post-seeding or day 0 of the differentiation protocol. (Figure 1a).

Following three-day culture on plates coated with basement membrane extract (BME), Geltrex, including a 24 hr treatment with 1% or 2% DMSO, and an untreated negative control we observed a slight increasing trend in the number of cells in the G1 phase correlated with an increase in DMSO concentration used. The percentage of cells in the G0/G1 phase was quantified by distinguishing cells with double amount of DNA to determine the phase of cell cycle (Figure 1b). We found that there was a substantial decrease in the overall number of cells with DMSO treatment (Figure 1c). To exclude the possibility of DMSO affecting proliferation, we assessed if the cells remaining in colonies were still proliferating. Immunofluorescent images of hiPSC colonies depicted localization of Ki67 protein expression, a marker of cell proliferation. Ki67 expression was observed in cells of all colonies untreated and treated with DMSO (Figure 1d), and this was further supported by flow cytometry data that showed a non-significant decrease in the percentage of cells expressing Ki67 with the increase of DMSO (Figure 1e). This suggests other reasons for reduced cell number, such as growth arrest, cell detachment or cell death.

Pluripotency was assessed following DMSO treatment. Immunofluorescent images of hiPSC colonies showed OCT4, and SOX2 protein expression throughout the hiPSC colonies (Figure 1l). Flow cytometry results for SSEA3, SSEA4, TRA-1-60, and TRA-1-81 (Figure 1 h-k) showed an overall high expression of characteristic pluripotent cell surface markers for all conditions. A significant decrease in protein expression of critical pluripotent transcription factors OCT4 and SOX2 (Figure 1 f-g) was observed following 2% DMSO treatment. We also measured gene expression of pluripotency transcription factors SOX2, c-myc, OCT4 and NANOG by qPCR analysis (Figure 1 m-p). SOX2 and c-myc remained stable (Figure 1 m-n). Interestingly, OCT4 expression decreased slightly (Figure 1 o) and expression of the non-critical pluripotent transcription factor, NANOG, decreased significantly (Figure 1 p). Other genes indicative of mesoderm differentiation (Brachy, T) was not affected by DMSO treatment (Figure 1 q). A gene associated with naive pluripotent human stem cells (Rex1) was found to be slightly upregulated with DMSO treatment but not significantly, (Figure 1 r), whereas gene expression of FGF5, a gene associated with primed pluripotency in mouse embryonic stem cells, was upregulated in cells treated with DMSO (Figure 1s). This analysis underscores the pluripotent state and differentiation potential of the cultured hiPSCs following treatment with DMSO based on gene and protein expression of key pluripotent markers.

### 3.2 Changes in hiPSC colony morphology following DMSO treatment

The impact of cell morphology was analysed upon DMSO exposure. Results shown in Figure 2 provide a parametric analysis of hiPSC colony morphology at the beginning (0 hr) and at the end (24 hrs) of the exposure to DMSO (Figure 2a). The LUMC-GFP+ hiPSCs pluripotency OCT4 and SOX2 (Figure S2a) was confirmed and with levels comparable to non-GFP LUMC line.

Fluorescence images of LUMC-GFP+colonies were taken at 20 min intervals over the course of the 24 hr DMSO treatment to identify the morphological changes in real-time (video S1-S3). A comparative analysis was performed on images acquired throughout 24 hr DMSO treatment (Figure 2b-e) using consistent settings on NIS software. The number of colonies was quantified in each image. The DMSO treatment resulted in an increase in the overall number of colonies for the start of differentiation as less colonies merged in wells treated with 1-2% DMSO compared to the non-treated control over the course of 24 hr (Figure 2b, S2a, Video S1a-c). This behavior was also observed for LUMCs imaged under brightfield (Video S2a-c). Quantification of fluorescent imaging of LUMC-GFP+ colonies revealed that 24 hr DMSO treatment leads to increase in colony circularity (Figure 2c, S2b) and convexity (Figure 2d), known to be specific morphological features of pluripotent stem cell colonies. Furthermore, the perimeter of the colonies was also assessed, which decreased with increasing DMSO treatment (Figure 2e, S2c). At 24 hr timepoint there was an overall statistical difference in all measurements between different conditions (Figure S1a-c). These measurements offered insights that clear morphological alterations to hiPSC colonies were observed following the DMSO treatment, which varied with the concentrations tested.

### 3.3 hiPSC structural rearrangement and integrin binding

With the changes previously reported on the colony morphology, a deeper investigation was performed on hiPSC cytoskeletal organization, as well as gene expression analysis related to integrins, and cell adhesion molecules.

Immunofluorescence images of the F-actin cytoskeleton, stained with Phalloidin (cyan), highlighted a gradual change at the cellular level and in the structural organisation of the colony (Figure 3a). Notably, there was a lack of pronounced cell protrusions, indicative of focal adhesions, on the periphery of the colonies when comparing colonies exposed to DMSO with non-treated conditions. Hence, the gene expression of critical components involved in integrin-mediated signalling and focal adhesions in stem cells were further analysed. Gene expression analysis revealed no change in E-Cadherin (CDH1) (Figure 3b), and upregulation of cell-cell adhesion molecule N-cadherin (CDH2) (Figure 3c) following the DMSO treatment. Focal adhesion kinase (PTK2), known to interact with integrin-mediated adhesion complexes and Vimentin were also downregulated with DMSO treatment (Figure 3d,e). The integrins (α5, α6, αν, β1, and β5), abundantly expressed on hiPSC and known to play important roles in binding induced hiPSCs to basement membrane components including collagen, fibronectin, laminin, and vitronectin were also analysed. The gene expression of most integrins remained unchanged with exception to both β1 and β5 integrins which was downregulated with DMSO treatment (Figure 3f-j).

We also observed morphological changes in colonies present in DMSO treated cultures, although more abundantly in 2% DMSO culture; small, condensed, circular colonies were observed (Figure S3a). These small colonies show lumen forming in the center (Figure 3m). Additionally, the lumen-forming protein podocalyxin was expressed at the top and centre of each colony (Figure 3k, m).

These findings collectively suggest that DMSO treatment significantly influences the cytoskeletal organization of hiPSCs and gene expression of integrin, cell adhesion molecules and adhesion sites on the periphery of cell colonies.

### 3.4 Metanephric mesenchyme nephron progenitor differentiation

The main aim of this study was to investigate the effects of DMSO treatment on hiPSC in differentiation efficiency, specifically in becoming nephron progenitors.

In figure 4 the impact of the DMSO treatment on the differentiation of NPCs was assessed. Cells were analysed at day 9 of the kidney organoid differentiation protocol, as depicted following the developmental timeline up to the stage when nephron progenitors emerge (Figure 4a). Immunofluorescence images showed hiPSCs successfully differentiated towards nephron progenitors presenting abundant expression of MM markers SIX2 (green) and PAX2 (red), when treated with 1% DMSO (Figure 4b,c). The percentage of MM nephron progenitors represented by SIX2+ cells in culture was quantitatively assessed using flow cytometry, providing insight into the efficiency of differentiation. The population of SIX2+ cells in non-treated hiPSCs reached 69%, treated with 1% DMSO reached on average 92%, and treated with 2% DMSO reached 83% (Figure 4d). Additionally, the number of cells per cm^2^ following dissociation was manually counted using a haemocytometer. Non-treated hiPSCs resulted on average in 6 × 10^5^cells/cm^2^, treated with 1% DMSO results in 12 × 10^5^ cells/cm^2^, and treated with 2% DMSO resulted in 7 × 10^5^ cells/cm^2^ (Figure 4e). The assessment of the MM progenitor differentiation was also confirmed with the LUMC-GFP+ expressing line, on day 9 of the kidney organoid differentiation and these also expressed the markers for nephron progenitors SIX2 and PAX2 (Figure S2b);

This comprehensive analysis underscores the successful differentiation of hiPSCs into nephron progenitors, where treatment of hiPSC with 1% DMSO leads to highest number of renal progenitors (identified as SIX2+ cells).

### 3.5 Kidney organoid differentiation

To assess if the DMSO treatment affects nephrogenesis, nephron progenitors (SIX2+ cells) were further differentiated and the efficiency of these to become kidney organoids containing developing nephron structures was acessed.

Results presented in Figure 5 showed the final stages of kidney organoid differentiation on day 21. A schematic outlines the 11 days differentiation post MM progenitors of kidney organoid culture (Figure 5a). The number of individual organoids developing in each well of a 96-well plate was manually counted for each condition. The percentage of wells with successfully differentiated kidney organoids for each condition is represented in Figure 5b-d. The results showed that 38% of the wells containing cells not treated with DMSO contained kidney organoids with characteristic tubular structure (Figure S1b) while 78% and 50% of wells contained kidney organoids for cells treated with 1% and 2% DMSO, respectively. A similar trend was observed under brightfield in 6 well plate format (Figure S1c). Furthermore, the degree of successful differentiation was again observed for the different hiPSCs lines investigated, with a clear advantage when 1% DMSO was used (Figure S1d). Total protein content results (Figure 5e) showed the protein signal correlated with the number of wells with no. of kidney organoid structures present.

The fluorescent signal for key proteins characteristic of different nephron sections was measured and normalised against total protein. Overall, the amount of podocalyxin (glomerulus) signal did not vary across conditions (Figure 5f). The signal for megalin (proximal tubule) increased in wells treated with higher concentraion DMSO (Figure 5g). A significant increase was measured for signal of GATA3 (distal tubule/collecting duct) in wells treated with 2% DMSO compared to non-treated control (Figure 5h). Immunofluorescent images of kidney organoids showed the structural arrangement of the developing nephrons for each condition and expression of characteristic proteins found in the different segments of the nephron. The kidney organoids differentiated in each condition showed expression of Podocalyxin (PODXL) for the podocytes in the glomerulus, lotus tetragonolobus lectin (LTL) for brush boarder of the proximal tubule, and E-cadherin located in the epithelium of the developing distal tubule and collecting duct (Figure 5i, S1e), as well as nephrin located at the slit diaphragm of the glomerulus, functional transporter protein megalin in the proximal tubule, and GATA3 located in the epithelium of the distal tubule and collecting duct (Figure 5j, S1e) in each of the respective tubular structures. The LUMC-GFP+ line on day 21 of kidney organoid differentiation also showed that the tubular structures emerging expressed nephron markers, namely Megalin and GATA3 (Figure S2c).

This analysis highlights the impact exposure of hiPSCs to DMSO has on the downstream differentiation of kidney organoids. The protein quantification presented some significant changes only in the expression of, the distal tubule/collecting duct marker GATA3, between tested conditions. We also found the treatment with 1% DMSO presented the highest rate of organoid formation and the lowest rate of failed differentiations.

## 4. Discussion

### 4.1 Changes Pluripotency and differentiation potential

This study aimed to investigate how DMSO treatment affects hiPSC differentiation, particularly for MM NPCs and kidney organoids. We examined the impact of a 24-hour DMSO treatment on hiPSCs. Previous research has identified cell surface markers like SSEA3, SSEA4, TRA-1-81, and TRA-1-60 as indicators of pluripotency [24, 25]. Our results showed that DMSO treatment did not alter the expression of these markers, suggesting that the pluripotent state was maintained at this level of analysis.

When examining transcription factors essential for maintaining pluripotency, such as SOX2 and OCT4, we found that OCT4 protein expression was downregulated following DMSO treatment while SOX2 gene expression remained stable. This observation aligns with previous studies that have shown DMSO can influence OCT4 expression during the differentiation of hepatic cells [26], suggesting that DMSO may modulate pluripotency at the transcriptional level without fully disrupting the gene expression profile necessary for maintaining hiPSC identity. Another transcription factor, NANOG, is normally expressed in pluripotent cells but primarily plays a role in reprogramming [27]. NANOG is associated with the high proliferation rate characteristic of pluripotent stem cells. Previous studies have shown that NANOG overexpression is linked to the rapid progression from the G1 to S phase of the cell cycle through the downregulation of the cell cycle inhibitor p27 [28, 29], further underscoring its role in maintaining a proliferative state. Interestingly, our study found that DMSO treatment resulted in a significant reduction in NANOG expression and a concomitant decrease in cell numbers after 24 hours. However, we did not observe much reduction in the expression of proliferation marker, Ki67, at a protein level. Our findings align with the known effects of DMSO on the cell cycle, where DMSO has been reported to increase the population of pluripotent stem cells in the G1 phase through regulation of genes associated with inhibition of the PI3K/Akt signalling pathway [17]. Inhibition of this pathway has been associated with downregulation of pluripotency markers and upregulation of lineage-specific genes, indicating a shift from a pluripotent state towards differentiation [30]. Studies have also shown inhibition of PI3K activity decreased expression of NANOG [31]. NANOG was previously found to be downregulated in cells transitioning through the primitive streak to form mesoderm and definitive endoderm [32]. We also measured the gene expression of mesoderm marker Brachy (T) but found no significant change in expression following DMSO treatment.

We also explored the hypothesis that DMSO treatment may push hiPSCs into a more “primed state” of pluripotency. The transition from naive to primed pluripotency involves molecular changes, including the expression of specific markers and transcription factors associated with the epiblast stage in mouse embryonic stem cells. Our study also found similar patterns in gene expression with as downregulation of NANOG [33], and upregulation of FGF5 [34]. N-cadherin also becomes detectable as cells exit the pluripotent state, and PSCs primed for neural and mesodermal differentiation show a shift from E-cadherin to N-cadherin expression [35]. Our findings showed that DMSO-treated hiPSCs exhibited upregulation N-cadherin, consistent with a shift toward a more “primed state” of pluripotency. N-Cadherin gene expression increased but no change for E-Cadherin was observed, indicating that this Cadherin switch had not yet occurred. As upregulation of N-Cadherin is also associated with epithelial to mesenchyme transition (EMT) we decided to investigate if the other EMT markers were upregulated with DMSO, but found that EMT marker Vimentin was in fact downregulated indicating more stable hiPSC colonies [36].

Taken together, these results indicate that while DMSO-treated hiPSCs maintain surface marker expression characteristic of pluripotency, there are significant changes at the transcriptional level, particularly involving key pluripotency factors such as SOX2, OCT4, and NANOG. These changes may signal the beginning of a transition to a different state of pluripotency, driven by the modulation of cell cycle progression and alterations in pluripotency-related transcription factors. Our study provides new insights into the dynamic effects of DMSO on hiPSCs, highlighting its potential role in influencing the balance between pluripotency maintenance and differentiation. Further investigation using a broader panel of molecular markers associated with primed and naive pluripotency could provide deeper insights into how DMSO influences hiPSC states and differentiation potential.

### 4.2 Changes in hiPSC colony morphology

DMSO-treated hiPSCs exhibited distinct morphological changes, prompting us to measure these alterations and correlate them with established knowledge about the relationship between hiPSC colony structure and pluripotency. Previous research has shown that compact colonies with tight cellular associations are characteristic of successful hiPSC conversion [1], and smaller colonies tend to have higher differentiation potential due to the greater proportion of cells at the colony edges [37]. This is further amplified by the increased number of individual colonies present in cultures treated with 1-2% DMSO.

Our results showed that colonies treated with DMSO presented a reduced perimeter, suggesting they are better candidates for differentiation. Additionally, DMSO treatment led to increased colony convexity and circularity, indicating less variability in colony morphology indicative of more stable hiPSCs [36]. Phalloidin staining revealed a more rounded cytoskeletal organization with fewer thin actin stress fibers at the colony edges, indicating possible alterations in mechanical forces and biophysical interactions within the colonies, potentially influencing their state of pluripotency and differentiation potential [38].

A notable observation was the emergence of small, round colonies with a lumen forming in the centre following 2% DMSO treatment consistent with previous reports [39]. We noticed this particular shape resembled epiblast spheroids [40]. These spheroids are reported to be primed for differentiation, particularly toward the mesoderm lineage [41]. They are characterised by formation of lumens with apicobasal polarity markers like PODXL, ZO-1, and β-catenin. Our study observed similar morphological features in DMSO-treated hiPSCs, suggesting that DMSO may induce a transition towards more “primed state” of pluripotency, similar to epiblast spheroids.

### 4.3 hiPSC structural rearrangement and interaction with BME

Cell protrusions indicative of focal adhesions on the periphery of hiPSC colony edges appear to be affected by DMSO treatment, leading us to investigate the effects on integrin expression and focal adhesion dynamics. Integrins, particularly α5, α6, αv, β1, and β5, play a crucial role in stem cell-matrix interactions and significantly influence human embryonic stem cell survival and differentiation [42]. Our analysis revealed that hiPSCs grown on BME, Geltrex - an extra-cellular matrix derived from Engelbreth-Holm-Swarm mouse sarcomas, expressed high levels of these integrins, consistent with previous reports. Following DMSO treatment, integrin expression levels remained stable, with a notable decrease in integrin β5 expression, potentially impacting cell attachment and proliferation [43]. Integrin interaction with extra-cellular matrix substrates activate focal adhesion kinase (FAK) and AKT signalling, which are critical for suppressing cytoskeletal contraction and preventing apoptosis [44, 45]. DMSO-treated hiPSCs showed slight downregulation of FAK expression, along with a loss of adhesion sites and a decrease in cell number, suggesting potential effects on cell proliferation or survival [17].These findings highlight the impact DMSO treatment has on hiPSC cell attachment and interaction with BME, which can explain the reason for the reduction of hiPSC survival following DMSO treatment.

### 4.4 Metanephric mesenchyme nephron progenitor differentiation

In the second part of our study, we evaluated if the treatment of hiPSCs with DMSO had a downstream effect on the generation of MM progenitors, essential for kidney organoid differentiation. It is known that DMSO enhances differentiation of cells towards the three germ layers [17-19]. However, the influence of DMSO on the generation of MM progenitors and kidney organoids has not been reported. Hence, we evaluated the differentiation protocol efficiency by measuring the SIX2+ cells, a crucial MM NPC marker [46]. Previous studies demonstrated that it is critical to obtain 80-90% SIX2+ cells [11, 20] for successful downstream nephrogenesis. We quantified the difference in SIX2+ cells populations on day 9 of differentiation protocol and found that, on average, 1% DMSO had highest percentage of MM induction with 92% SIX2+ population, followed by highest cells retrieval rate, likely due to higher proliferation of MM NPCs in the culture. Treating hiPSC with 1% DMSO was an effective way for creating a higher number of NPCs from a single batch differentiation culture. This would be particularly beneficial when generating many uniform 3D kidney organoids are needed for downstream applications (e.g., toxicity screening).

### 4.5 Mature kidney organoid nephron differentiation

We also aimed to evaluate the success rate of kidney organoid generation in 96 well plate format, and structural composition of the resulting nephron development. Results showed that 1% DMSO treatment had the highest success rate of kidney organoid differentiation, followed by 2% DMSO, and lastly, non-treated hiPSCs. Kidney organoids development for each condition showed similar structures with the expression of characteristic markers for different sections of the nephron: glomerulus (Podocalyxin and Nephrin), proximal tubule (LTL and Megalin), and distal tubule/collecting duct (E-cadherin and GATA3) [8, 10, 11, 21]. We observed that kidney organoids derived from hiPSCs treated with DMSO appear to have slightly higher protein expression of markers in the proximal tubule (megalin) and significantly higher expression of distal tubule/collecting duct (GATA3), whereas expression of markers for the glomerulus (podocalyxin) remained unchanged. These results suggest that treating hiPSCs with DMSO and subsequent differentiation following our optimised protocol resulting in kidney organoids with a higher number of tubular structures. GATA3 is not exclusively expressed in the distal tubule but also in the collecting duct arising from the ureteric bud lineage [10]. Therefore, it may be plausible that including DMSO treatment of hiPSCs leads to more cells differentiating into anterior intermediate mesoderm and subsequently differentiating towards the collecting duct. More analysis is needed to determine exactly which lineages emerge during the differentiation process [47]. These findings show that DMSO treatment enhances the differentiation of kidney organoids with developing nephron structures, particularly the distal tubule and collecting duct. By refining protocols with simple interventions, this work improves the efficiency of kidney organoid generation which can have significant impact on biomedical research and drug development.

### 5. Conclusion and future perspectives

We found that hiPSCs treated with 1-2% DMSO continue to express critical transcription genes and proteins essential for maintaining pluripotency. However, genes associated with a ‘primed’ pluripotent state and post-implantation epiblast were affected by DMSO treatment. Furthermore, we found that treating hiPSC colonies with DMSO resulted in more individual colonies before commencing differentiation and more circular colonies with less protrusions concurrent with focal adhesions on the colony edges, a key morphological parameter used for phenotype classification of hPSC colony differentiation potential. With 1-2% DMSO treatment we also observed the emergence of colonies with shape, apico-basal polarity, and lumen formation in the centre is indicative of epiblast spheroid morphology.

Following kidney organoid differentiation, we showed an overall number of cells and percentage population expressing the critical metanephric mesenchyme marker, SIX2, in DMSO-treated hiPSCs compared to non-treated controls. 1% DMSO was found to give rise to the highest percentage of SIX2+ cells during differentiation and the highest success rate for fully mature organoids on day 21 of the differentiation protocol. Kidney organoids cultured until day 21 of differentiation protocol expressed characteristic markers for different segments of the developing nephron for each condition, with 2% DMSO-treated hiPSCs showing significantly higher expression of GATA3+ cells than non-treated control. The application of DMSO enhances organoid differentiation, increasing the yield of tubular kidney organoids and improving scalability for future toxicity screening and regenerative medicine applications, where a high number of organoids are often required. Future studies are needed to fully elucidate the alterations of gene expression from pluripotency and through the different stages of differentiation with a deeper analysis on a single cell level. Focus should be given to the first few days of differentiation to determine the rate of which cells treated with DMSO progress toward the primitive streak. This would potentially allow deeper understanding on how DMSO affects differentiation toward mesoderm.

## Declarations

### Ethical Approval

LUMC0031iCTRL08: LUMCi004-C (RRID: CVCL_ZA01) (https://hpscreg.eu/cell-line/LUMCi004-C) were generated from exfoliated renal epithelial cells with commercially available episomal plasmid. Cells were isolated from human samples with informed consent and ethics approval in compliance with the relevant laws. The human iPSC lines HUMIMC101: TISSUi001-A (RRID:CVCL_WU56) (https://hpscreg.eu/cell-line/TISSUi001-A) and HUMIMC107: TISSUi007-A (RRID:CVCL_WU62) (https://hpscreg.eu/cell-line/TISSUi007-A) were derived from peripheral blood mononuclear cells. Human blood samples were donated with informed consent and ethics approval in compliance with the relevant laws.

### Consent to Participate

Approval and consent for iPSCs-lines has been secured by the respective entities that generated the cells.

### Consent to Publish

All authors have given their consent for the publication of the manuscript.

### Authors Contributions

Conception and design of the study: HK, CM. Data acquisition: HK, AR, AS, EE. Analysis and interpretation of data: HK, AR, AS, EE, CM. Drafting and revision of the manuscript: HK, AR, AS, EE, PW, LM, CM. All authors have approved the final manuscript.

### Availability of data and materials

The data supporting the findings of this study are available from the corresponding author upon reasonable request.

