## Supplementary material for "Dimethyl sulfoxide primes induced pluripotent stem cells for more efficient nephron progenitor and kidney organoid differentiation"

### 1 Supplementary Information

Table S1: List of antibodies for flow cytometry, immunofluorescence images and in-cell western.

| Antibody name | Host | Dilution | Manufacturer | Catalog # |
|---|---|---|---|---|
| TRA-1-81 | Mouse | 1:10 | BD bioscience | 560161 |
| TRA-1-60 | Mouse | 1:10 | BD bioscience | 560193 |
| SSEA3 | Rat | 1:40 | BD bioscience | 561145 |
| SSEA4 | Mouse | 1:40 | BD bioscience | 560219 |
| SOX2-PE | human | 1:100 | Miltenyi Biotech | 130-121-053 |
| OCT3/4-AF647 | Mouse | 1:100 | BD Bioscience | 560329 |
| SOX2 | Mouse | 1:200 | Abcam | ab79351 |
| OCT4 | Rabbit | 1:250 | Abcam | ab200834 |
| Podocalyxin | Goat | 1:200 | R&D systems | AF1658 |
| Beta-catenin | Rabbit | 1:200 | Abcam | ab2365 |
| ZO-1 | Mouse | 1:200 | Invitrogen | 33-9100 |
| Phallodin | n/a | 1:200 | Invitrogen | 10125092 |
| Six homeobox 2 | Rabbit | 1:300 | Proteintech | 11562-1-AP |
| PAX2 | Goat | 1:100 | R&D systems | AF3364 |
| Lotus tetragonolobus lectin | n/a | 1:300 | Vector laboratories | FL-1321-2 |
| E-cadherin | Mouse | 1:150 | BD bioscience | 610181 |
| Megalin | Mouse | 1:300 | R&D Systems | MAB9578-100 |
| Nephrin | Sheep | 1:300 | R&D systems | AF4269 |
| GATA3 | Rabbit | 1:500 | Cell signalling | 5852S |
| 4′,6-diamidino-2-phenylindole | n/a | 2ng/mL | Sigma-Aldrich | 32670 |
| Donkey anti-goat AF568 | Donkey | 1:1000 | Invitrogen | 10463972 |
| Donkey anti-mouse AF647 | Donkey | 1:1000 | Invitrogen | 15980296 |
| Donkey anti-sheep AF568 | Donkey | 1:1000 | Invitrogen | A21099 |
| Donkey anti-rabbit AF488 | Donkey | 1:1000 | Invitrogen | 10424752 |
| Donkey anti-goat AF647 | Donkey | 1:1000 | Invitrogen | 10493402 |
| Goat anti-mouse AF647 | Goat | 1:500 | Invitrogen | A-21240 |
| Goat anti-mouse AF568 | Goat | 1:1000 | Invitrogen | A-21245 |
| Goat anti-mouse IRDye 800CW | Goat | 1:500 | LICOR | 926-32210 |
| Goat anti-rabbit IRDye 800CW | Goat | 1:500 | LICOR | 926-32211 |
| Donkey anti-goat IRDye 800CW | Donkey | 1:500 | LICOR | 926-32214 |

Table S2: List of primers used for qPCR.

| Gene ID | Forward | Reverse |
|---|---|---|
| ATP5PB | TTTCATACAGGGCAGCCACA | AAGCCCAGTTCCGAGTACAT |
| SOX2 | GCTTAGCCTCGTCGATGAAC | AACCCCAAGATGCACAACTC |
| OCT4 | GGTTCTCGATACTGGTTCGC | GTGGAGGAAGCTGACAACAA |
| c-myc | CCTGGTGCTCCATGAGGAGAC | CAGACTCTGACCTTTTGCCAGG |
| Nanog | ACCAGTCCCAAAGGCAAACA | TCTGCTGGAGGCTGAGGTAT |
| Brachy (T) | AGGTACCCAACCCTGAGGA | GCAGGTGAGTTGTCAGAATAGGT |
| Rex1 | CGCAATCGCTTGTCCTCAGAGT | GCTCTCAACGAACGCTTTCCCA |
| FGF5 | GGAATACGAGGAGTTTTCAGCAAC | CTCCCTGAACTTGCAGTCATCTG |
| Vimentin | GCCGAAAACACCCTGCAATC | TCCTGGATTTCCTCTTCGTGG |
| CDH2 | CCTCCAGAGTTTACTGCCATGAC | GTAGGATCTCCGCCACTGATTC |
| PTK2 | GCCTTATGACGAAATGCTGGGC | CCTGTCTTCTGGACTCCATCCT |
| CD151 | GGAGAACCTGAAGGACACCATG | CAGTCCTGTGAGTTGTTGCTGC |
| ITGa5 | GCCGATTCACATCGCTCTCAAC | GTCTTCTCCACAGTCCAGCAAG |
| ITGa6 | CGAAACCAAGGTTCTGAGCCCA | CTTGGATCTCCACTGAGGCAGT |
| ITGaV | AGGAGAAGGTGCCTACGAAGCT | GCACAGGAAAGTCTTGCTAAGGC |
| ITGb1 | GGATTCTCCAGAAGGTGGTTTCG | TGCCACCAAGTTTCCCATCTCC |
| ITGb5 | GCCTTTCTGTGAGTGCGACAAC | CCGATGTAACCTGCATGGCACT |
| PODXL | AACCCGGCCCAAGATAAGTG | TTGGCAGGGAGCTTAGTGTG |
| β-Catenin | CACAAGCAGAGTGCTGAAGGTG | GATTCCTGAGAGTCCAAAGACAG |

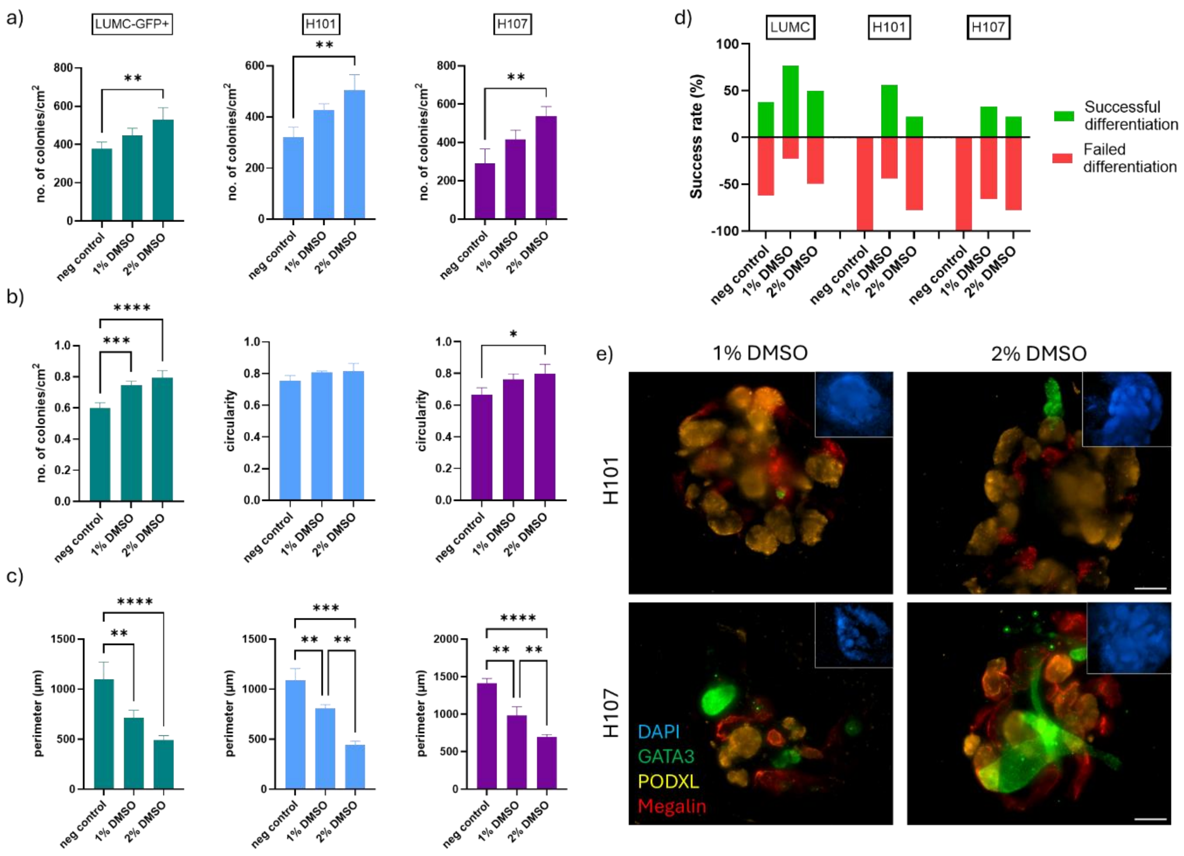


**Figure S1 – Parametric analysis of three different hiPSCs treated with 0-2% DMSO for 24hr prior to kidney organoid differentiation.** Nuclei of cells in colonies were imaged LUMC-GFP, H101 and H107 and used to a) count the number of cell colonies, b) measure the perimeter of cell colonies, and c) cell colony circularity; LUMC-GFP+ (N=3), H101 (n=3) and H107 (n=3). On day 21 of kidney organoid differentiation the d) success rate of kidney organoid differentiation for the three different hiPSC lines was assessed in each well of 96 well plate (LUMC n=60, H101 n=9 and H107 n=9). e) Immunofluorescent images confirming expression of nephron markers in kidney organoid structures from differentiation cultures using H101 n=1 and H107 n=1; Podocalyxin (yellow), Megalin (red) and GATA3 (green), scale bar – 100 µm.

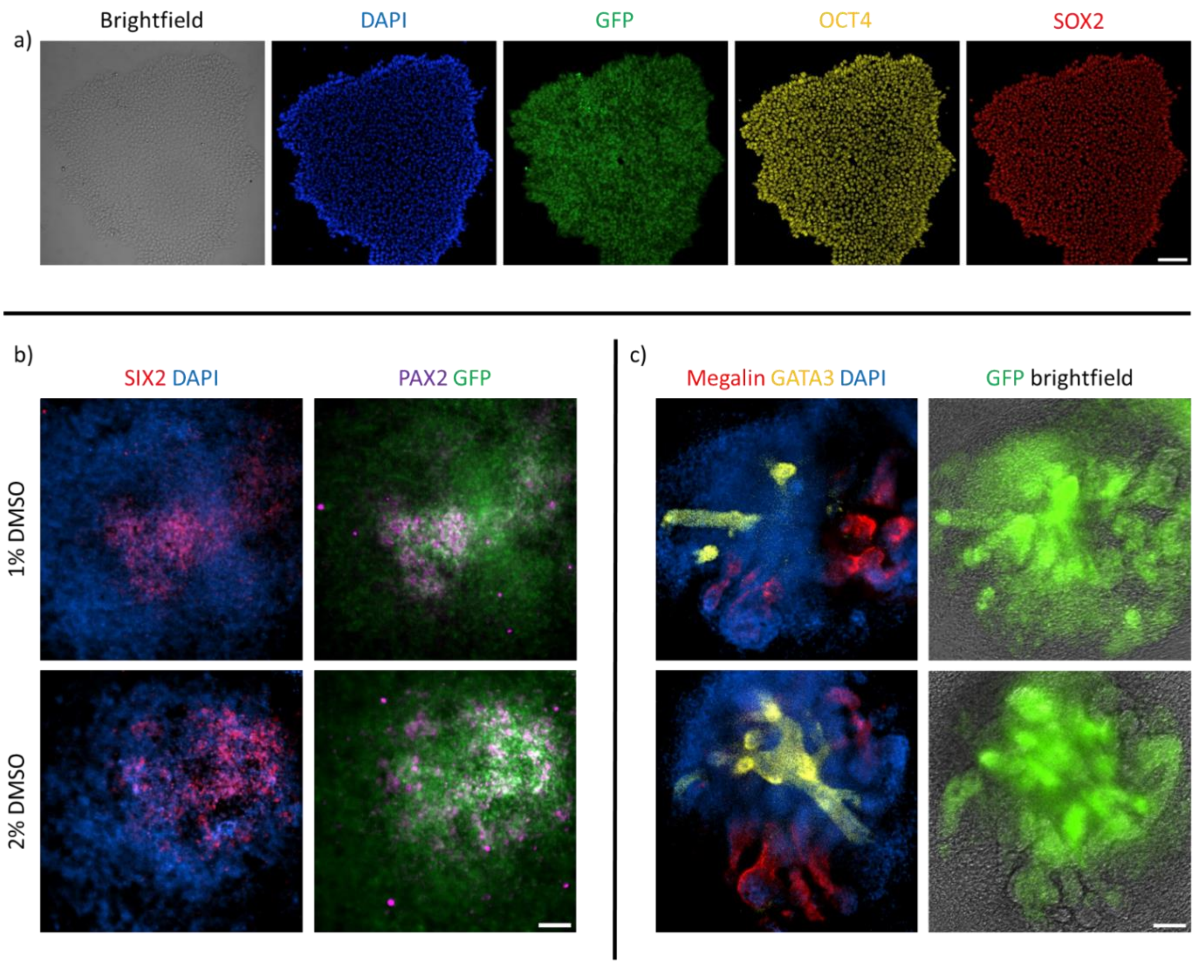


**Figure S2 –** Immunofluorescent images of a) LUMC-GFP+ hiPSCs colony (green) stained for markers of pluripotency OCT4 (yellow) and SOX2 (red), and nuclear DAPI (blue) staining. LUMC-GFP+ hiPSCs differentiated towards b) MM nephron progenitors expressing SIX2 (red), PAX2 (magenta), and nuclear DAPI (blue) staining on day 9 of kidney organoid differentiation, and c) kidney organoids expressing nephron markers Megalin (red), GATA3 (yellow), and nuclear DAPI (blue) staining on day 21 of kidney organoid differentiation (n=1). Scale bar – 100 µm.

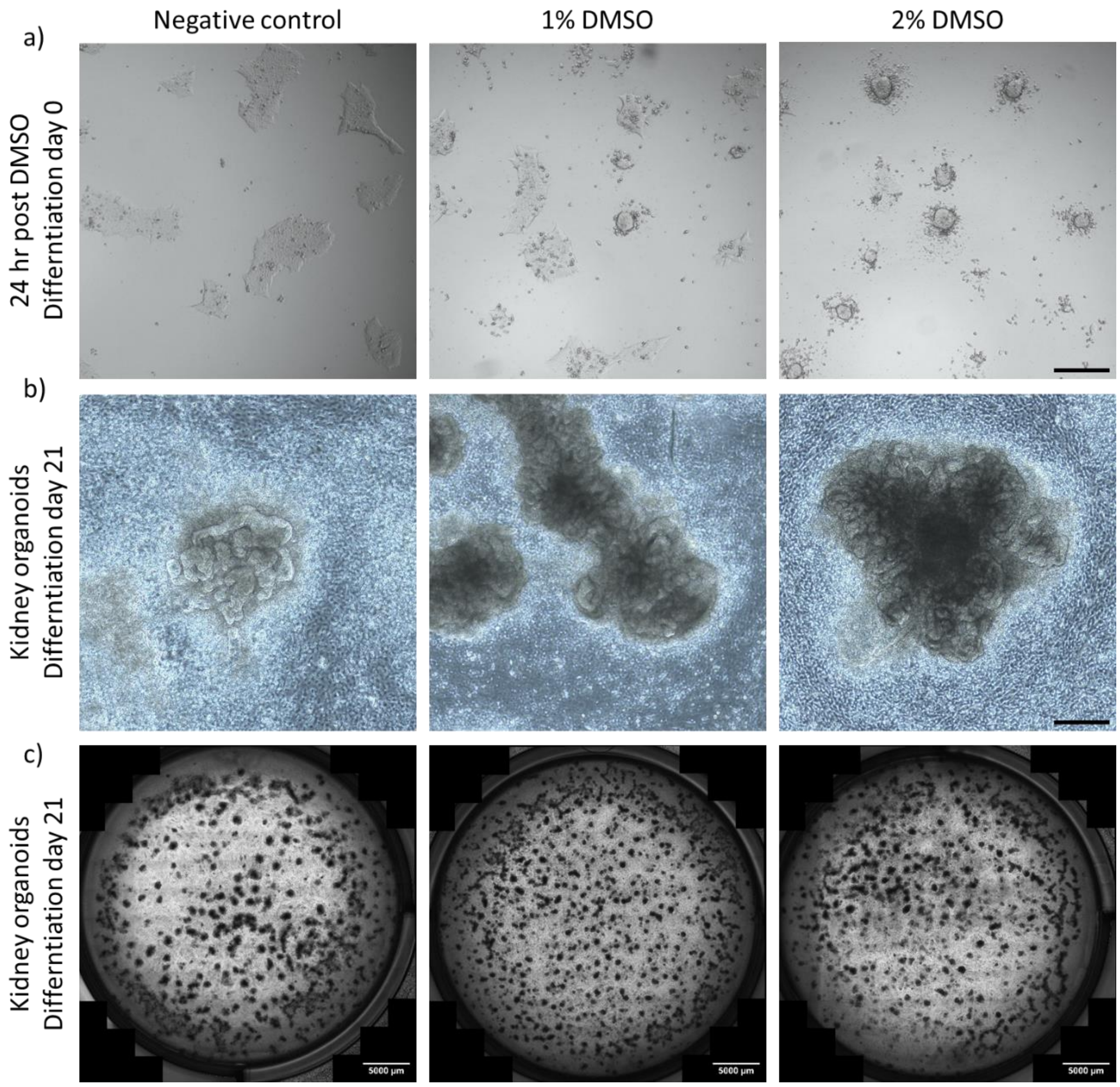


**Figure S3 –** Brightfield images of a) LUMC hiPSCs treated with DMSO for 24hr, scale bar - 100 µm, b) kidney organoids differentiated until day 21, scale bar - 100 µm, and c) stitched image of full well in a 6 well plate with kidney organoids differentiated until day 21, scale bar - 500 µm.

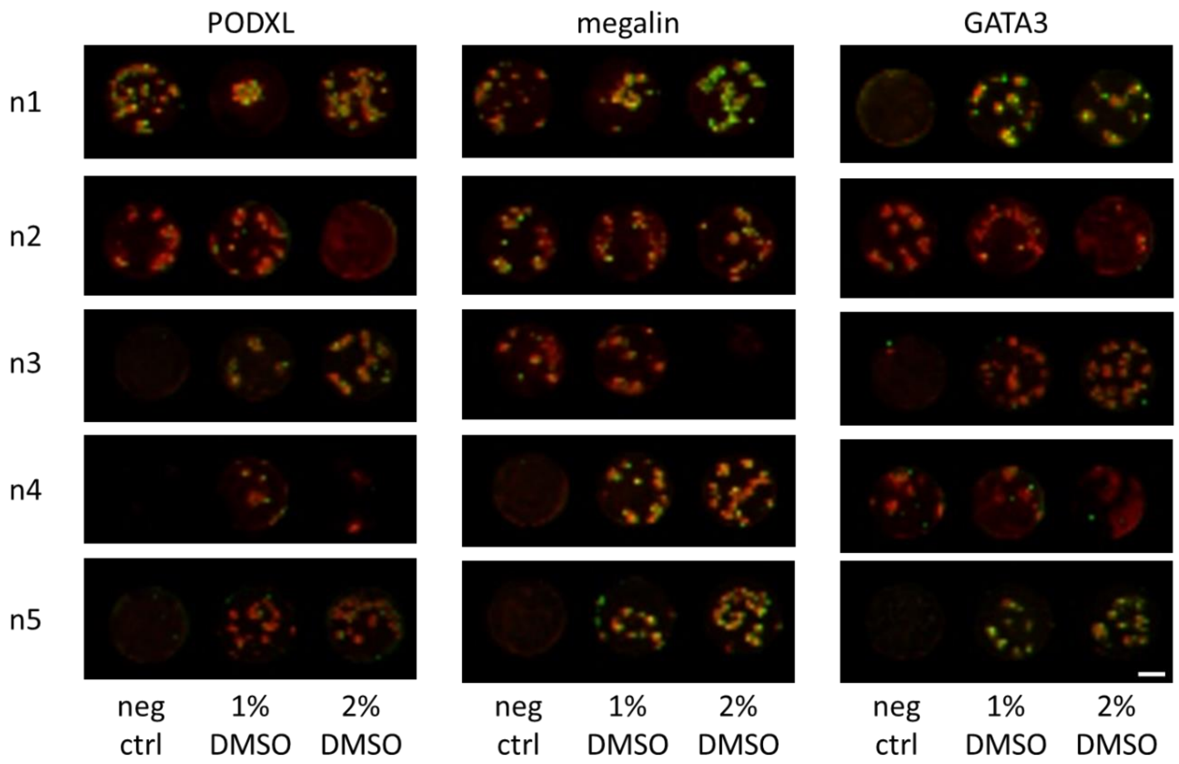


**Figure S4 –** LICOR NIR images of wells used for quantifying total protein - CellTag700 (red) and Podocalyxin, Megalin and GATA3 (green). Scale bar – 200 µm

**Video S1a –** Timelapse video of LUMC-GFP+ colonies under brightfield of non-treated control well (negative control). Single images taken every 20 min over 24 hr.

**Video S1b –** Timelapse video of LUMC-GFP+ colonies under brightfield treated with 1% DMSO. Single images taken every 20 min over 24 hr.

**Video S1c –** Timelapse video of LUMC-GFP+ colonies under brightfield treated with 2% DMSO. Single images taken every 20 min over 24 hr.

**Video S2a –** Timelapse video of LUMC colonies under brightfield non-treated control well (negative control). Single images taken every 20 min over 24 hr.

**Video S2b –** Timelapse video of LUMC colonies under brightfield treated with 1% DMSO. Single images taken every 20 min over 24 hr.

**Video S2c –** Timelapse video of LUMC colonies under brightfield treated with 2% DMSO. Single images taken every 20 min over 24 hr.
